# Non-specific recognition of histone modifications by H3K9bhb antibody

**DOI:** 10.1101/2023.04.12.536655

**Authors:** Takeshi Tsusaka, Juan A. Oses-Prieto, Christina Lee, Brian C. DeFelice, Alma L. Burlingame, Emily Goldberg

**Affiliations:** Department of Physiology, University of California, San Francisco, San Francisco, CA 94158, USA; Department of Pharmaceutical Chemistry, University of California, San Francisco, San Francisco, CA 94158, USA; Department of Molecular and Cell Biology, University of California, Berkeley, Berkeley, CA 94720; Chan-Zuckerberg Biohub, San Francisco, CA 94158 USA

**Keywords:** ketone, β-hydroxybutyrate, β-hydroxybutyrylation, Kbhb, post-translational modification, antibody specificity

## Abstract

Ketone bodies are short chain fatty acids produced in the liver during periods of limited glucose availability that provide an alternative source of energy for the brain, heart, and skeletal muscle. Beyond this classical metabolic role, β-hydroxybutyrate (BHB), is gaining recognition as a pleiotropic signaling molecule. Lysine β-hydroxybutyrylation (Kbhb) is a newly discovered post-translational modification in which BHB is covalently attached to lysine ε-amino groups. This novel protein adduct is metabolically sensitive, dependent on BHB concentration, and found on proteins in multiple intracellular compartments, including the mitochondria and nucleus. Therefore, Kbhb is hypothesized to be an important component of ketone body-regulated physiology. Kbhb on histones is proposed to be an epigenetic regulator, which links metabolic alterations to gene expression. However, we found that the widely used antibody against the β-hydroxybutyrylated lysine 9 on histone H3 (H3K9bhb) also recognizes other modification(s), which are increased by deacetylation inhibition and include likely acetylations. Therefore, caution must be used when interpreting gene regulation data acquired with the H3K9bhb antibody.

## Introduction

Post-translational modifications (PTMs) are important regulators of cell biology through their ability to modify protein function, structure, and location. Importantly, many PTMs are directly regulated by metabolism. For example, histone acetylation is achieved by transfer of an acetyl group from acetyl-CoA ^1–4^), a central metabolic intermediate required for both ATP production and energy storage pathways. Through this mechanism, nutrient availability and cellular energy status can control gene expression ^5,6^).

While acetylation is a highly prevalent histone PTM, many other short chain fatty acids can also form covalent adducts with lysine, including lactylation ^7^, butyrylation ^8^, crotonylation ^9^, and propionylation ^8^. Recently, the ketone body β-hydroxybutyrate (BHB) was reported as a novel PTM on lysine residues, β-hydroxybutyrylation, an adduct referred to as Kbhb ^10^. BHB is the most abundant circulating ketone body, with baseline concentrations ~0.1-0.2mM in rodents and humans. BHB is produced by hepatocytes as an alternative energy source for the brain, heart, and skeletal muscle and is induced during periods of starvation or very-low-carbohydrate consumption, such that BHB levels increase >10-fold ^11,12^. This classical function of BHB underscores its essential physiological importance. In mammalian cell culture and mouse models, Kbhb abundance correlates with increasing concentrations of BHB ^10,13^. Therefore, there is great interest in understanding the significance of Kbhb and how this metabolically-sensitive PTM might mediate metabolic adaptations to negative energy balance.

Like many PTMs, Kbhb was originally discovered on histones ^10^. Many of the histone Kbhb sites are the same residues that can also be acetylated. Among the most studied Kbhb residues is histone 3 lysine 9 (H3K9) based on its association with active gene expression when it is acetylated (H3K9ac) at promoter regions ^14^. To date, several studies have used H3K9bhb chromatin immunoprecipitation (ChIP) assays to identify BHB-regulated genes in the context of BHB-treatment, starvation, or ketogenic diet ^10,15–20^).

Here we report our examination of the H3K9bhb antibody specificity. In the course of our studies, we observed unexpected induction of H3K9bhb antibody signal in cells not exposed to BHB. These data suggested the H3K9bhb antibody used in prior publications might not be specific to its intended target. Using mass spectrometry, we confirmed that H3K9bhb is enriched only in cells treated with BHB, and that western blot signal intensity does not reflect H3K9bhb abundance. Therefore, caution must be used when interpreting data obtained with the H3K9bhb antibody.

## Results

### Unexpected pattern of H3K9bhb expression

To test the specificity of Kbhb antibodies, we treated HEK293T cells and immortalized murine embryonic fibroblasts (iMEFs) with BHB, the structurally similar butyrate (Figure 1A), or the histone deacetylase inhibitor Trichostatin A (TSA) (Figure 1B). As expected, western blot analysis with a pan-Kbhb antibody showed many β-hydroxybutyrylated proteins of a wide range of molecular weights in BHB-treated cells, but not with butyrate or TSA treatment. We observed a similar pattern with a site-specific Kbhb antibody that recognizes H4K8bhb. Surprisingly, both monoclonal and polyclonal antibodies that detect H3K9bhb did not follow this pattern. BHB treatment increased the H3K9bhb signals, as expected. However, the H3K9bhb signal in cells treated with butyrate or TSA was comparable to or exceeded that of BHB-treated cells. These results were evident regardless of BHB concentration (Figure S1). After carefully reexamining the literature, we found this unexpected result was also previously observed ^21–23^, although it was not investigated further. Untargeted metabolomics confirmed that butyrate treatment does not increase intracellular BHB levels (Figure 1C). Collectively, these data suggested the H3K9bhb antibody might also recognize other histone modifications.

**Figure 1.**
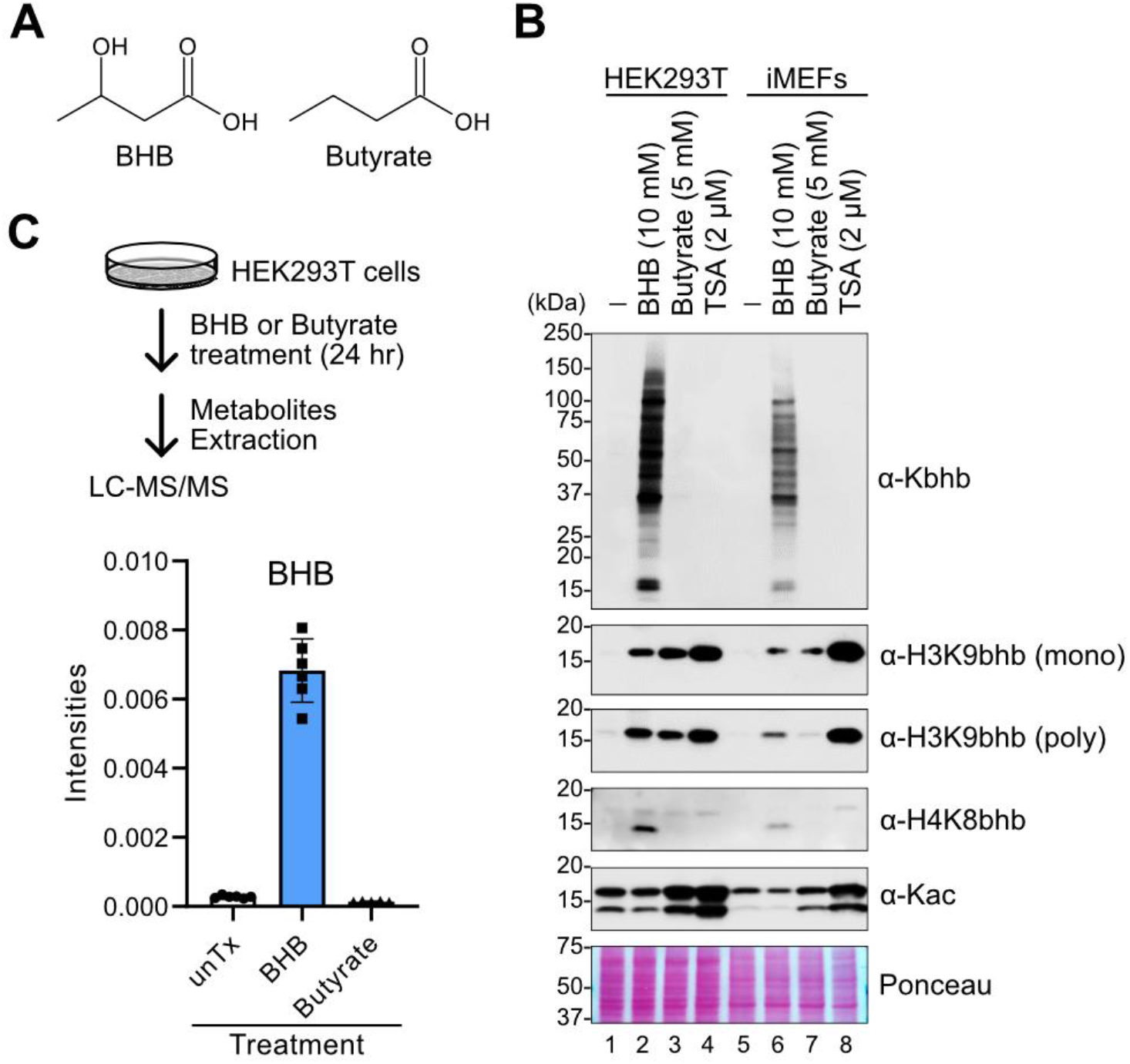
Irregular band patterns of H3K9bhb antibodies. **(A)** Structures of BHB and structurally similar butyrate. **(B)** Western blots of HEK293T cells and iMEFs treated with either BHB, Butyrate, or TSA for 24 hours. Butyrate and TSA are known deacetylation inhibitors. Ponceau S staining was used to confirm the equivalent loading of proteins. mono: monoclonal, poly: polyclonal. Data are representative of at least 3 independent experiments. **(C)** Upper: schematic of experimental workflow. HEK293T cells were treated with 5 mM BHB or 5 mM Butyrate. At 24 hours after the treatment, cellular metabolites were extracted and subjected to untargeted metabolomics by LC-MS/MS. unTx indicates untreated HEK293T cells. Lower: relative intensities of BHB in the metabolome of each treatment condition.

### Assessment of H3K9 modifications by mass spectrometry

Next, we directly tested the possibility that the H3K9bhb antibody might recognize alternative histone modifications. We treated HEK293T cells with BHB or butyrate and used the H3K9bhb antibody for immunoprecipitation (IP) (Figure 2A). Western blot analysis and ponceau staining confirmed the successful enrichment of H3K9bhb after IP in the BHB-treated sample (Figure 2B, lanes 4 vs 7). Mass spectrometry analysis confirmed the presence of H3 peptides that harbor BHB adducts on lysine 9 (Figures 2C, S2). As expected, Kbhb-containing H3 peptides were enriched in the BHB-treated sample, accounting for 27/183, or 13.99% (Table S1). In contrast, we only identified 2/113, or 1.74%, Kbhb-containing peptides in the butyrate-treated sample. A similar trend was also observed for the H3K9-containing peptide such as “K_9_STGGKAPR”: 3/14, or 21.43% for the BHB-treated sample and 0/24 for the butyrate-treated sample. Given that butyrate inhibits protein deacetylation ^24^, we queried the mass spectrometry data for acetylated peptides and found a strong enrichment of H3K9ac-containing peptides in the butyrate-treated sample as compared to the BHB-treated sample (Figure 2D, Table S1). We expanded our analysis and also found other prevalent PTMs in our H3K9bhb pulldown samples, including butyrylation, mono-, di-, and tri-methylation, or propionylation (which includes both endogenous PTM and chemical derivatization as indicated in Figure 2A). While these data do not conclusively pinpoint which modifications are recognized by the H3K9bhb antibody, this mismatch between H3K9bhb abundance in Figure 1B and the lack of H3K9bhb-containing peptides in Figure 2D indicate the antibody is not specific to H3K9bhb.

**Figure 2.**
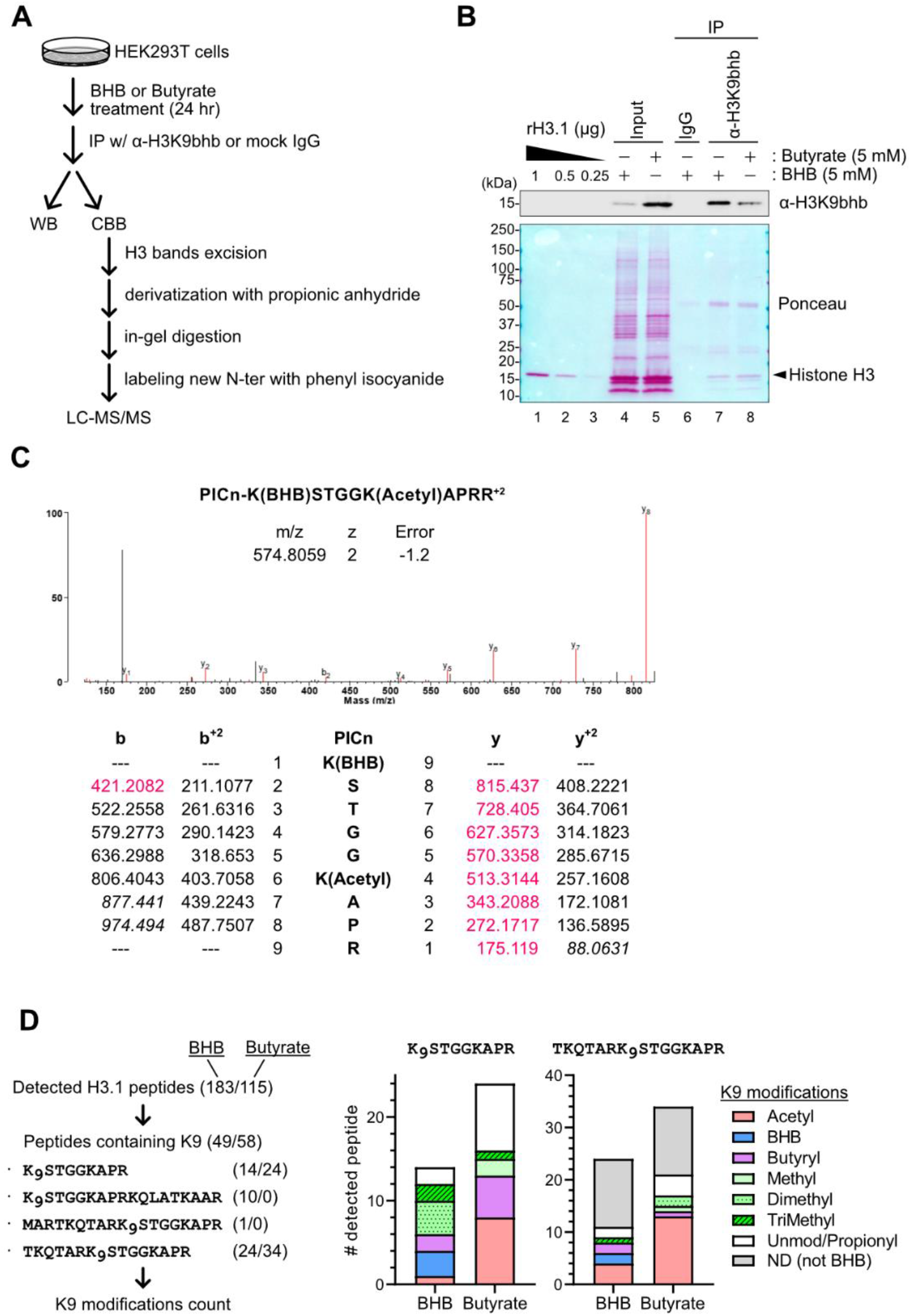
Absence of H3K9bhb in butyrate-treated HEK293T cells. **(A)** Schematic of experimental workflow. HEK293T cells were treated with 5 mM BHB or 5 mM butyrate for 24 hours. Cell lysates were subjected to immunoprecipitation with α-H3K9bhb antibody or control normal rabbit IgG. The immunoprecipitants were used for western blot analysis and SDS-PAGE, followed by Coomassie brilliant blue (CBB) staining and LC-MS/MS analysis. **(B)** Western blot of input and H3K9bhb IP fractions of HEK293T treated with BHB or butyrate for 24 hours. Ponceau S staining was used to confirm the equivalent loading of proteins and H3 enrichment. **(C)** Representative tandem mass (MS/MS) spectra of the K9-BHB-ylated and K14-acetylated “PICnKSTGGKAPRR” peptides from BHB-treated samples. The detected y ions and b ions are highlighted in pink. **(D)** Left: a strategy of the downstream analysis. Numbers in parentheses refer to the number of detected peptides in BHB or butyrate-treated samples. Right: detected peptide numbers for each modification at the K9 position in the indicated peptides. ND: not determined.

## Discussion

Histone modification antibodies are notorious for specificity problems due to structural similarities and possible binding site access limitations due to neighboring site modifications^25–27^. Our data demonstrate conclusively that the only commercially available H3K9bhb antibody recognizes additional modifications, likely including acetylation, that undermines the reliability of this reagent for ChIP experiments to assess H3K9bhb-regulated gene expression. Of note, we expected to recover more H3K9bhb-containing peptides with our pulldown strategy (Figure 2). We speculate this could be an indication of low prevalence of H3K9bhb in cells, even after treatment with high concentrations of BHB. We hope that by publicly reporting our data, new reagents can be developed with improved specificity so future studies can reexamine the possible importance of H3K9bhb. In addition, prior datasets using H3K9bhb (summarized in Table 1) should be interpreted with caution and the possible detection of H3K9ac or other PTMs should be considered.

**Table 1.** Summary of previous studies that used H3K9bhb antibody. This table contains the prior publications that used the PTM Biolabs H3K9bhb antibodies, what sample type was analyzed, and what readouts were used. WB: western blot.

| Study | Cells/Tissues Analyzed | H3K9bhb Application |
| --- | --- | --- |
| Xie et al., 2016 | mouse liver, HEK293 | ChIP-seq, WB |
| Chen et al., 2017 | mouse hypothalamus, primary neurons | WB |
| Nishitani et al., 2018 | 3T3-L1 | ChIP-qPCR |
| Chriett et al., 2019 | HEK293, L6 myotubes, HMEC-1 | WB |
| Zhang et al., 2020 | mouse CD8+ T cells | WB, ChIP-qPCR |
| Zhang et al., 2019b | HEK293T | Dot blot, WB |
| Huang et al., 2021 | HCT116, MEF, HeLa | WB |
| Terranova et al., 2021 | mouse small intestine crypt cells | ChIP-seq, WB |
| Zhang et al., 2021 | hepatocellular carcinoma | WB |
| Sangalli et al., 2022 | dairy cow cells/tissues | WB |
| Zheng et al., 2022 | mouse pancreas, Bx-PC3 cells | ChIP-Seq, CUT&Tag, WB |
| Cornuti et al., 2023 | occipital cortex | ChIP-seq, WB |
| Tang et al., 2023 | mouse CD8+ T cells | WB, ChIP-qPCR |

## Acknowledgements

We thank members of the Goldberg lab for comments and intellectual discussion. We also thank Hiten Madhani (UCSF) for intellectual discussion and manuscript feedback, Anna Molofsky’s lab (UCSF) for sharing HEK293T cells, and Yoshihiro Ishikawa (UCSF) for sharing iMEFs. We also thank Catharine Bosio, Benjamin Schwarz, and Eric Bohrnsen (NIAID/NIH Rocky Mountain Labs) for feedback and independent metabolomics validation. This work was funded by NIH/NIA (R00AG058801 to ELG, a pilot and feasibility award from the UCSF Liver Center P30DK026743 to ELG), the Chan Zuckerberg Biohub, the Sandler Program for Breakthrough Biomedical Research, which is partially funded by the Sandler Foundation (to ELG), JSPS Overseas Research Fellowships (to TT), and the Dr. Miriam and Sheldon G. Adelson Medical Research Foundation (AMRF, to Dr. A. L. Burlingame, Director of the Mass Spectrometry Resource Center at UCSF).

## Author Contributions

ELG and TT conceptualized project, designed experiments, and prepared the manuscript. TT performed experiments and analyzed data. JAOP performed mass spectrometry sample prep, acquisition, and data analysis, and this was overseen by ALB. CL performed experiments. BDF performed untargeted metabolomics sample prep, acquisition, and analysis. BS and EB performed metabolomics sample acquisition and data analysis, and this was overseen by CB. All authors read, commented on, and approved the manuscript.

## STAR Methods

### KEY RESOURCES TABLE

| REAGENT or RESOURCE | SOURCE | IDENTIFIER |
| --- | --- | --- |
| Antibodies |  |  |
| anti-Kbhb | PTM Biolabs | Cat# PTM-1201RM |
| anti-H3K9bhb (mono) | PTM Biolabs | Cat# PTM-1250 |
| anti-H3K9bhb (poly) | PTM Biolabs | Cat# PTM-1250RM |
| anti-H4K8bhb | PTM Biolabs | Cat# PTM-1253RM |
| anti-Kac | Cell Signaling Technology | Cat# 9814S; RRID: AB_10544700 |
| Normal rabbit IgG | Cell Signaling Technology | Cat# 2749 |
| Goat anti-Rabbit IgG HRP | Thermo Scientific | Cat# 31460 |
| Rabbit TrueBlot®: Anti-Rabbit IgG HRP | ROCKLAND | Cat# 18-8816-31 |
| Chemicals, peptides, and recombinant proteins |  |  |
| DL-β-Hydroxybutyric acid sodium salt (BHB) | Sigma-Aldrich | H6501 |
| Sodium butyrate | Sigma-Aldrich | B5887 |
| Trichostatin A (TSA) | Sigma-Aldrich | T8552 |
| RIPA buffer | Thermo Scientific | 89901 |
| Protease inhibitor cocktail | Thermo Scientific | A32965 |
| Dynabeads™ M-280 Sheep Anti-Rabbit IgG | Invitrogen | 11203D |
| DMEM (Dulbecco's Modified Eagle's Medium) | Gibco | 10569044 |
| DPBS (Dulbecco's Phosphate Buffered Saline) | Gibco | 14190250 |
| Antibiotic-Antimycotic (100X) | Gibco | 15240062 |
| Critical commercial assays |  |  |
| DC Protein Assay | BioRad | 5000116 |
| Experimental models: Cell lines |  |  |
| HEK293T | N/A | N/A |
| iMEFs | ATCC | CRL-1503 |
| Software and algorithms |  |  |
| Prism (v9.5.1.) | Graphpad | https://www.graphpad.com/ |
| ProteinProspector (v 6.4.0) | (Guan et al., 2011) | https://sargent.ucsf.edu/prospector/mshome.htm |

#### Experimental model and subject details

##### Cell culture

HEK293T cells and iMEFs (immortalized mouse embryonic fibroblasts) were cultured in DMEM (Gibco) supplemented with 10% fetal bovine serum (FBS) and 1x antibiotic-antimycotic (Gibco) in a 37°C incubator with 5% CO2. Cells were treated with BHB, butyrate, and TSA at the indicated concentrations in each figure for 24 hours.

#### Method details

##### Western blot analysis

Cells were harvested after trypsinization and quenching with medium, washed once with PBS, and lysed with RIPA buffer (Thermo Scientific) supplemented with protease and phosphatase inhibitors (Thermo Scientific). The cell lysates were sonicated using a Branson Sonifier 450 until the viscosity of lysates disappears and centrifuged at 14,000 rpm (17,968 x1*g*) for 5 mins. The supernatants were subjected to the DC Protein Assay Kit (BioRad Laboratories). SDS-PAGE samples were prepared by mixing the supernatants and 4x Laemmli buffer/10% beta-mercaptoethanol (BioRad Laboratories) and followed by boiling at 95°C for 5 mins. Equivalent amounts of protein were resolved by SDS-PAGE and then transferred to a nitrocellulose membrane using the Trans-Blot Turbo Transfer system (BioRad Laboratories). Membranes were stained with Ponceaus S, blocked with 5% milk/TBS-T (Tris Buffered Saline/0.1% Tween-20) for 30-60 mins, and incubated with primary antibodies, followed by secondary HRP-conjugated antibodies. Blots were developed using a chemiluminescent substrate (SuperSignal™ West Pico PLUS or SuperSignal™ West Femto Maximum Sensitivity Substrate; Thermo Scientific) and imaged with an Azure 300 (azure biosystems).

##### Sample preparation for untargeted metabolomic analysis

Cells were seeded at the dense of 6.0×10^5 cells/well of a 6-well dish. Six wells were used per condition. After one day of culture, BHB or butyrate at 5 mM was directly added to the culture medium. At 24 hours after treatment, cells were washed with 0.9% NaCl at room temperature, harvested by scraping, centrifuged at 1,500 rpm to remove the supernatant, and stored as a pellet at −80°C until further processing. Cell pellets were resuspended with 225 μL of ice-cold methanol and mixed with 750 μL of ice-cold methyl butyl ether (MTBE). After vortexing for 10 seconds and shaking for 6 mins at 4°C, 188 μL of distilled water was added to the samples. Samples were additionally vortexed for 20 seconds and centrifuged for 2 min at 14,000x*g*. The bottom phase was transferred to a new tube, dried, subjected to lipid cleanup, resuspended with 20 μL 3:2 v/v acetonitrile:water, and used for LC-MS/MS.

##### LC-MS/MS using the HILIC (hydrophilic interaction liquid chromatography) method

We performed untargeted metabolomics using the HILIC method to detect polar metabolites including BHB as described previously ^28^. Samples were injected onto a Waters Acquity UPLC BEH Amide column (150 mm length × 2.1 mm id; 1.7 μm particle size) with an additional Waters Acquity VanGuard BEH Amide pre-column (5 mm × 2.1 mm id; 1.7 μm particle size) maintained at 45°C coupled to an Thermo Vanquish UPLC. The mobile phases were prepared with 10 mM ammonium formate and 0.125% formic acid in either 100% LC-MS grade water for mobile phase (A) or 95:5 v/v acetonitrile:water for mobile phase (B). Gradient elution was performed from 100% (B) at 0–2 min to 70% (B) at 7.7 min, 40% (B) at 9.5 min, 30% (B) at 10.25 min, 100% (B) at 12.75 min, isocratic until 16.75 min with a column flow of 0.4 mL/min. Spectra were collected using a Thermo Q Exactive HF Hybrid Quadrupole-Orbitrap mass spectrometer in both positive and negative mode ionization (separate injections). Full MS-ddMS2 data was collected, an inclusion list was used to prioritize MS2 selection of metabolites from the in-house ‘local’ library, when additional scan bandwidth was available MS2 was collected in a data-dependent manner. Mass range was 60-900 mz, resolution was 60k (MS1) and 15k (MS2), centroid data was collected, loop count was 4, isolation window was 1.2 Da.

##### Immunoprecipitation (IP)

Cells were treated with BHB or butyrate at 5 mM. After 24 hours of treatment, approximately 5.0×10^7 cells per sample were harvested after trypsinization and quenching with medium, washed once with PBS, and lysed with PBS-T (PBS/1% Triton X-100) supplemented with protease and phosphatase inhibitors (Thermo Scientific). After incubating for 10 min on ice, the cell lysates were centrifuged at 10,000 rpm (9,168 x*g*) for 5 mins to remove supernatant (rough cytoplasmic fraction). The cell pellet was washed once with 1 mL PBS-T, snap-frozen by liquid nitrogen, and stored at −80°C until further processing. The frozen pellet was resuspended in 500 uL SDS-lysis buffer (50 mM Tris-HCl at pH7.5, 1% SDS) per 5.0×10^7 cells. The lysates were sonicated using a Branson Sonifier 450 and were subjected to the DC Protein Assay Kit (BioRad Laboratories). 7 mg/mL lysates were made by adding SDS-Lysis buffer and beta-mercaptoethanol at a final concentration of 0.5%, and were boiled at 95°C for 5 mins to denature proteins. The denatured proteins were diluted 1:10 with NP-40 lysis buffer (50 mM Tris-HCl at pH 7.5, 150 mM NaCl, 1 mM EDTA, 10% glycerol, 1% Nonidet P-40) to reduce SDS concentration. The approximately 0.7 mg/mL lysates (5 mL per sample) were incubated with antibody-conjugated anti-Rabbit IgG Dynabeads (Invitrogen, 11203D) overnight at 4 °C. The resins were then washed five times with the NP-40 lysis buffer and eluted by incubating them in 0.2% SDS-elution buffer (50 mM Tris-HCl at pH7.5, 0.2% SDS) at 60°C for 20 mins. Elutions were mixed with 4x Laemmli buffer/10% beta-mercaptoethanol (BioRad Laboratories) and boiled at 95°C for 5 mins. The 1/10 amount of samples was used for western blot analysis and the remainder was used for mass-spectrometry analysis.

##### Histone hybrid propionylation-phenyl isocyanate derivatization, in gel digestion and mass spectrometry analysis

Immunoprecipitated samples were loaded on SDS-PAGE and stained with Coomassie Brilliant Blue (CBB; GelCode Blue Stain; Thermo Scientific). A band corresponding to histone H3 was excised using a sharp and clean blade. H3 bands were excised, cut in 1mm^3^ pieces, and transferred to low protein binding 0.6 ml tubes. Proteins were digested in-gel with trypsin as described previously ^29^ and H3 in gel was subjected to a hybrid propionylation-phenyl isocyanate derivatization, adapting the method described by Maile ^30^ with the necessary modifications to processing in gel samples, as indicated next. Gel pieces were washed for 5 min with 100 μl 50% acetonitrile (MeCN) in 100 mM triethylammonium bicarbonate (TEAB) pH 8.5, with shaking. The supernatant was removed, and samples rehydrated by adding 100 μl 100 mM TEAB pH 8.5 and shaking for 5 min. The supernatant was removed. The washes with 50% MeCN in 100 mM TEAB and then 100 mM TEAB were repeated 3 more times, until the Coomassie staining was removed. Then we added to the gel pieces 80 μl 100 mM TEAB pH 8.5, to reach a total volume including gel around 100 μl; then added 10 μl of propionic anhydride 1/100 in water, vortexing and incubating the samples for 5 min at room temperature. After that, we removed the supernatant, and repeated 2 more times the incubation with propionic anhydride, adding each time 80 μl 100 mM TEAB pH 8.5 plus 10 μl of propionic anhydride 1/100 in water. After removing the supernatant, we added 90 μl 100 mM TEAB pH 8.5 plus 10 μl of 80 mM hydroxylamine to quench excess reagent and incubated for 20 min at room temperature with shaking. After that, we removed the supernatant and washed for 5min with 100 μl 50% MeCN in 100 mM TEAB, then 3 times with 100 μl 70% MeCN in 100 mM TEAB and evaporated the gel pieces to total dryness in speed vac. Dry gel pieces were added 8 ul of a Trypsin solution (Promega) 10ng/μl in 100 mM TEAB. After incubating at room temperature for 10 min to allow complete absorption of the solution in the acrylamide pieces, we added 40 μl 100 mM TEAB and incubated with shaking at 37C, overnight. After that, we added 15 μl of a 1% v/v solution of phenyl isocyanate (PIC) in acetonitrile (Prepared fresh) to each sample, and incubated for 60 min at 37 °C. The reaction with PIC was repeated again. Then, the gel pieces were sonicated in bath for 5 min, and the supernatant recovered in a new set of 0.6 ml low protein binding vials. The gel pieces were extracted twice with 40 μl 50% MeCN 5% formic acid, for 30 min with shaking, and the supernatant recovered to the tubes used previously to transfer the aqueous supernatants. These extracts were evaporated in a speed vac, resuspended in 20 μl of 0.1% formic and peptides extracted with with C18 ziptips (Millipore). Eluted peptide mixtures were evaporated and resuspended in 0.1% formic acid and run onto a 2 μm 75μm x 50 cm PepMap RSLC C18 EasySpray column (Thermo Scientific) using 3-hour MeCN gradients (2–30% in 0.1% formic acid), coupled to a Orbitrap Exploris (Themo Scientific) for analysis in positive ion mode. MS spectra were acquired between 365 and 1400 m/z with a resolution of 120000. For each MS spectrum, multiply charged ions over the selected threshold (2E4) were selected for MSMS in cycles of 3 seconds with an isolation window of 1.6 m/z. Precursor ions were fragmented by HCD. Acquisition method specified an inclusion list for the m/z of ions corresponding to H3 K9-R17 peptides with either BHB or butyrate, plus propionyl, mono, di and trimethyl, acetyl, butyryl or BHB. A dynamic exclusion window was applied which prevented the same m/z from being selected for 30s after its acquisition.

For peptide and protein identification, peak lists were generated using PAVA in-house software ^31^. All generated peak lists were searched against the human subset of the SwissProt database (SwissProt.2019.07.31), using Protein Prospector ^32^ with the following parameters: Enzyme specificity was set as Arg-C, and up to 2 missed cleavages per peptide were allowed. N-acetylation of the N-terminus of the protein, loss of protein N-terminal methionine, pyroglutamate formation from of peptide N-terminal glutamines, oxidation of methionine, PIC derivatization of N termini (C7 N1 O1 H5), methylation and dimethylation of arginine, phosphorylation of serine, threonine or tyrosine, propionylation, acetylation, methylation, dimethylation, trimethylation, ubiquitination, butyrylation and β -hydroxybutyrylation (+86.0368) of lysine, were allowed as variable modifications, and up to 6 modifications per peptide were allowed. Mass tolerance was 5 ppm in MS and 30 ppm in MS/MS. The false positive rate was estimated by searching the data using a concatenated database which contains the original SwissProt database, as well as a version of each original entry where the sequence has been randomized. A 1% FDR was permitted at the protein and peptide level. As a downstream analysis, peptides corresponding to histone H3.1 were extracted and counted. Subquently, H3K9-containing peptides were extracted and counted. Each modification on the lysine 9 positions was manually counted. All the peptides information was given in Table S1.

