## Supplemental Figures for "Non-specific recognition of histone modifications by H3K9bhb antibody"

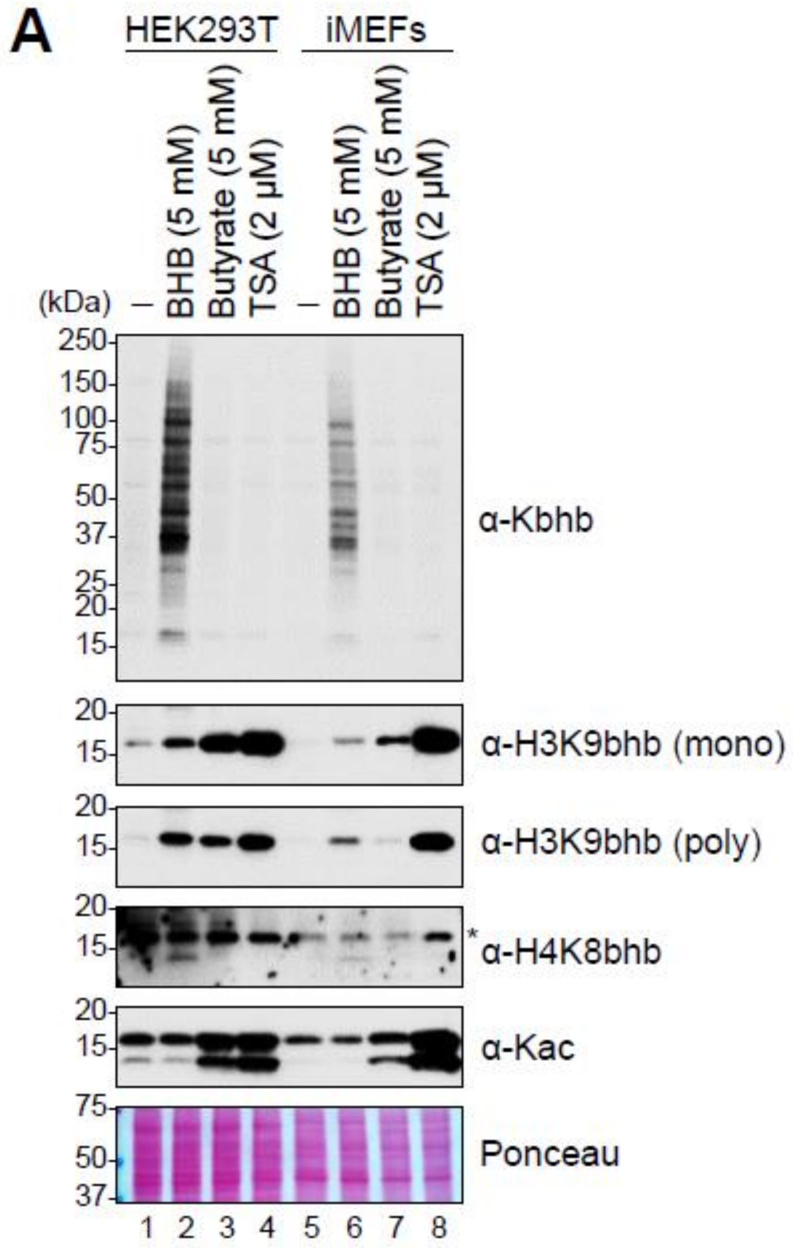


**Supplementary Figure 1, related to Figure 1. Reproducible observation of the unexpected signals when using anti-H3K9bhb antibodies.**

Western blots of HEK293T cells and iMEFs, as performed in Figure 1B, with the exception of using 5 mM BHB instead of 10 mM.

**A**

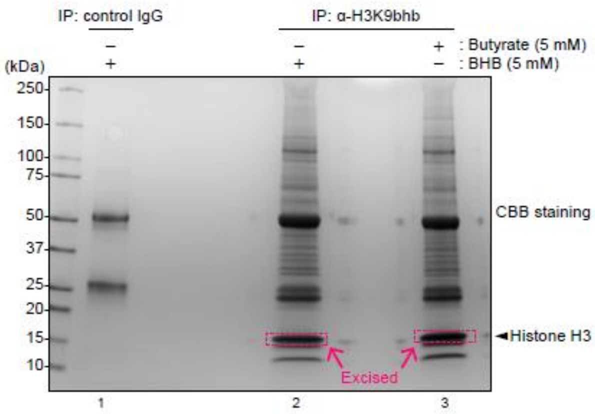


**B**

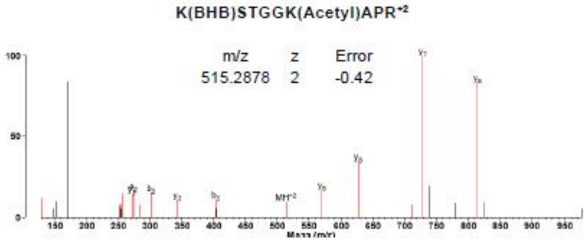


| b | b+2 | | | | y | y+2 |
|---|---|---|---|---|---|---|
| --- | --- | 1 | K(BHB) | 9 | --- | --- |
| 302.171 | 151.5892 | 2 | S | 8 | 815.437 | 408.2221 |
| 403.2187 | 202.113 | 3 | T | 7 | 728.405 | 364.7061 |
| 460.2402 | 230.6237 | 4 | G | 6 | 627.3573 | 314.1823 |
| 517.2617 | 259.1345 | 5 | G | 5 | 570.3358 | 285.6715 |
| 687.3672 | 344.1872 | 6 | K(Acetyl) | 4 | 513.3144 | 257.1608 |
| 758.4043 | 379.7058 | 7 | A | 3 | 343.2088 | 172.1081 |
| 855.4571 | 428.2322 | 8 | P | 2 | 272.1717 | 136.5895 |
| --- | --- | 9 | R | 1 | 175.119 | 88.0631 |

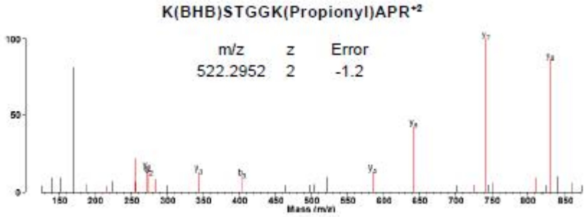


| b | b+2 | | | | y | y+2 |
|---|---|---|---|---|---|---|
| --- | --- | 1 | K(BHB) | 9 | --- | --- |
| 302.171 | 151.5892 | 2 | S | 8 | 829.4526 | 415.23 |
| 403.2187 | 202.113 | 3 | T | 7 | 742.4206 | 371.7139 |
| 460.2402 | 230.6237 | 4 | G | 6 | 641.3729 | 321.1901 |
| 517.2617 | 259.1345 | 5 | G | 5 | 584.3515 | 292.6794 |
| 701.3828 | 351.1951 | 6 | K(Propionyl) | 4 | 527.33 | 264.1686 |
| 772.4199 | 386.7136 | 7 | A | 3 | 343.2088 | 172.1081 |
| 869.4727 | 435.24 | 8 | P | 2 | 272.1717 | 136.5895 |
| --- | --- | 9 | R | 1 | 175.119 | 88.0631 |

**Supplementary Figure 2, related to Figure 2. LC-MS/MS analysis on histone H3 from BHB or Butyrate-treated cells**

**(A)** Coomassie brilliant blue (CBB) staining of IPed samples, as described in Figure 2A. H3 bands were excized and subjected to mass spectrometry analysis. **(B)** MS/MS spectra for the indicated peptides. The detected y ions and b ions are highlighted in pink.
